# Dispersal syndromes allow understanding but not predicting dispersal ability across the tree of life

**DOI:** 10.1101/2024.04.01.587575

**Authors:** Guillermo Fandos, Robert A. Robinson, Damaris Zurell

## Abstract

Dispersal is fundamental to many ecological and evolutionary processes, yet understanding the determinants and predictability of dispersal remains a crucial challenge. Dispersal syndromes, which describe patterns in the covariation of traits associated with dispersal, can help to gain a deeper understanding of the evolutionary dynamics of dispersal and its implications for range dynamics and ecosystem functioning in the face of global change. However, the extent to which such dispersal syndromes are generalizable across a large taxonomic scale has been hampered by low availability of standardized dispersal data across species. In this study, we used the most comprehensive and up-to-date empirical dispersal dataset for European birds to investigate the formation of dispersal syndromes and their effectiveness in predicting dispersal across species. We found body mass, feeding guild, and life-history traits shape dispersal syndromes in birds. Yet, only body mass and life history accurately predicted dispersal for unassessed species, while even multi-trait dispersal syndromes poorly predicted dispersal for phylogenetically distant species. These results shed light on the complex nature of bird dispersal and emphasize the need for refined approaches in elucidating the mechanisms and constraints underlying dispersal evolution. Our study highlights the importance of considering multiple factors and expanding empirical datasets to enhance our understanding of dispersal in avian populations.

## Introduction

Dispersal —the movement from a natal or breeding site to another breeding site— is a key ecological process that determines connectivity between populations and, by that, the population dynamics, gene flow, range dynamics, and local adaptations ^1–3^. It is increasingly recognized as a complex phenotypic process that emerges from trade-offs between selective pressures, environmental context, and covariation among traits ^4,5^. Dispersal syndromes reflect these patterns of covariation between morphological, physiological, or behavioral traits and dispersal ^6^ and provide critical insights into the evolution and causes of dispersal. Empirically measured dispersal data are scarce ^7,8^, and thus, dispersal syndromes have only been studied for a limited number of species ^9^ and in meta-analyses across different taxonomic groups rather than for entire clades. At the same time, dispersal information is crucial for making predictions of the species’ spatial dynamics under climate ^3,10,11^ and land-use change ^12–14^. Thus, a deeper understanding of dispersal syndromes is also key to improving our predictive capacity to anticipate future global change impacts on biodiversity.

Birds are one of the best-studied taxa, and indeed several studies have attempted to understand the covariation of dispersal ^15^ or dispersal proxies ^16,17^ with species traits. Yet, these previous studies have been limited to few species or by non-availability of standardized dispersal information and have not tested predictability of dispersal and thus the generality of the dispersal syndromes. From theoretical and empirical studies, we can expect that dispersal syndromes are multidimensional and that dispersal correlates with morphological, behavioral and life-history traits (reviewed in ^6^). In birds, morphological adaptations to dispersal are diverse, but variation in dispersal ability between species is found to positively covary with body size ^15^ or with any trait reducing movement costs, such as the wingspan or wing morphology ^16,17^. Similarly, behavior and physiology can strongly influence metabolism and, hence, the locomotor activity of a species ^18,19^. Functional and behavioral traits such as feeding guild ^20^, migratory behavior ^15^, or life-history traits (e.g., survival, age at maturity, and fecundity ￼) can also influence dispersal, for example, through variation in the behavioral strategies or spatial requirements for avoiding kin competition. Hitherto, relationships between dispersal and morphological, behavioral, and functional traits seem idiosyncratic among species and are rather noisy without robust empirical support, making identifying dispersal syndromes difficult ^22^. This has also been hampered by the inconsistencies in measurements and definitions of dispersal and the inability of most studies to detect long-distance dispersal events ^9,23^. Identifying the set of traits that shape short and long-distance dispersal syndromes can help us to elucidate the mechanistic determinants of dispersal, the constraints associated with movement, and, more generally, the trade-offs associated with dispersal evolution.

An open question is whether dispersal syndromes offer general and predictable insights into the dispersal process and can hence be used for gap-filling dispersal information for other species ^24^. Previously, simplified or theoretically derived dispersal syndromes have been used for this purpose ^25,26^ as empirical dispersal estimates are scarce ^7,8,15^, These studies have used species-level traits (body mass, wingspan or wing morphology, migratory behavior, life history) ^25–27^ and phylogenetic relatedness ^10^ to predict dispersal for diverse taxonomic groups. However, the full potential of these trait-based approaches to predict dispersal is currently limited by several conceptual and methodological factors. One limitation arises from the potential confusion between dispersal (the movement between consecutive breeding sites) and non-directed movement or migration (including nomadic or seasonal movements). Birds rely on highly adaptive traits to undertake seasonal migrations ^28,29^ that may not be identical to the traits relevant for natal/breeding dispersal. Furthermore, as the scarce empirical dispersal data have impeded the robust assessment of these dispersal inferences across many species, families, and orders, little is known about how generalizable or predictable dispersal syndromes are between closely to distantly related species. Recent developments of open-access databases on bird dispersal ^8^, phylogeny ^30^, and species traits ^31,32^ are now providing an unparalleled chance to understand the strength and directionality of dispersal syndromes and, consequently, the reliability of using a trait-based approach to infer dispersal ability in bird species with missing data.

Here, building on new dispersal estimates from ^8^, we explore the existence of dispersal syndromes in European bird species. Birds are ecologically diverse and show a range and variability of traits across species, which allows robust comparisons across the group and facilitates comprehensive analyses and investigations. Also, they are the only group for which standardized dispersal estimates exist across many closely and more distantly related species. We first investigate systematic covariations between (natal and breeding) dispersal ability and a suite of traits, while controlling for how common ancestry influences this covariation. We include traits associated with demography, morphology, ecological specialization, and behaviors relevant to movement ^33^ and assess their overall importance for explaining interspecific variation in dispersal. Within this analysis, we also test for consistency of dispersal syndromes between median-distance (explorative or routine movement) versus long-distance (sporadic) dispersal. Finally, we apply a cross-validation statistical approach to determine how single-trait and multi-trait models can predict dispersal distances, both within and across Orders. This evaluation will provide valuable insights and recommendations for researchers seeking to fill gaps in bird dispersal estimates in cases where empirical evidence is limited or unavailable.

## Results

### Detection of dispersal syndromes

Information about median and long dispersal average estimates were available for 234 species, and natal and breeding dispersal estimates for 113 European breeding bird species ^8^. Focusing on two heavy-tailed kernels that best explain dispersal movement in European birds (the Half-Cauchy and the Weibull distributions) ^8^, we tested covariations between bird dispersal ability (median and long-distance) and a suite of morphological, ecological and biogeographic traits (Table S1): body size, Hand Wing Index (HWI), slow-fast continuum of life histories (from long-lived species with high adult survival representing the slower end to early maturing species with short generation times and high reproductive rates ^34^), habitat openness, feeding guild, breeding latitude, and migration tendency. Multi-trait phylogenetic regression models included, according to the ecological knowledge, interaction terms between body size and life history, diet and habitat openness ^15,22^. We found similar dispersal syndromes using the Weibull and Half-Cauchy distribution (Fig.1, Fig. S1). For simplicity, we show the results of the dispersal syndromes from the Weibull distribution only. In the case of average (breeding and/or natal) dispersal, the strongest predictors of median dispersal distances were body mass and its interaction with life history (z = 0.367 and z = −0.222, respectively; see Figs. 1-2). Body mass is traditionally used as an index for dispersal in vertebrates, with a larger size assumed to indicate greater dispersal ability (e.g.^15,26^. We found an interaction between body mass and life-history (Fig. 1; Fig. S5), where the slow-fast continuum positively affected dispersal, but only in small birds, while larger birds show a negative correlation. We also found that species breeding in lower latitudes and migrating larger distances have greater dispersal ability compared to species of higher latitudes or shorter-distance migrants (z = −0.160, and z= 0.144; Fig. 1). Although the effect of feeding guild was weak (z= 0.018), we found a significant interaction between body mass and diet, where larger birds that feed obligatorily on animals tend to have longer dispersal distance than larger birds that feed on plants (z= 0.094). Associations between dispersal with habitat openness and its interaction with body mass were weaker (z = - 0.056 and z = −0.023, respectively; see Fig. 1; Fig. S4), but still selected in the final model. Overall, we found similar patterns and robust results for the median and long-distance dispersal estimates, except for the effect of life history. Similarly, we did not find great differences between average, natal and breeding dispersal syndromes patterns (Fig. 2, Fig. S4), which are, in any case, moderately well correlated (r > 0.5) ^8^. The main exceptions were a stronger effect of diet on natal than breeding dispersal (z = 0.166 and z = 0.465; Figs. 1-2) and a weakly negative effect of HWI on natal dispersal (z = −0.136; Fig. S4).

**Figure 1.**
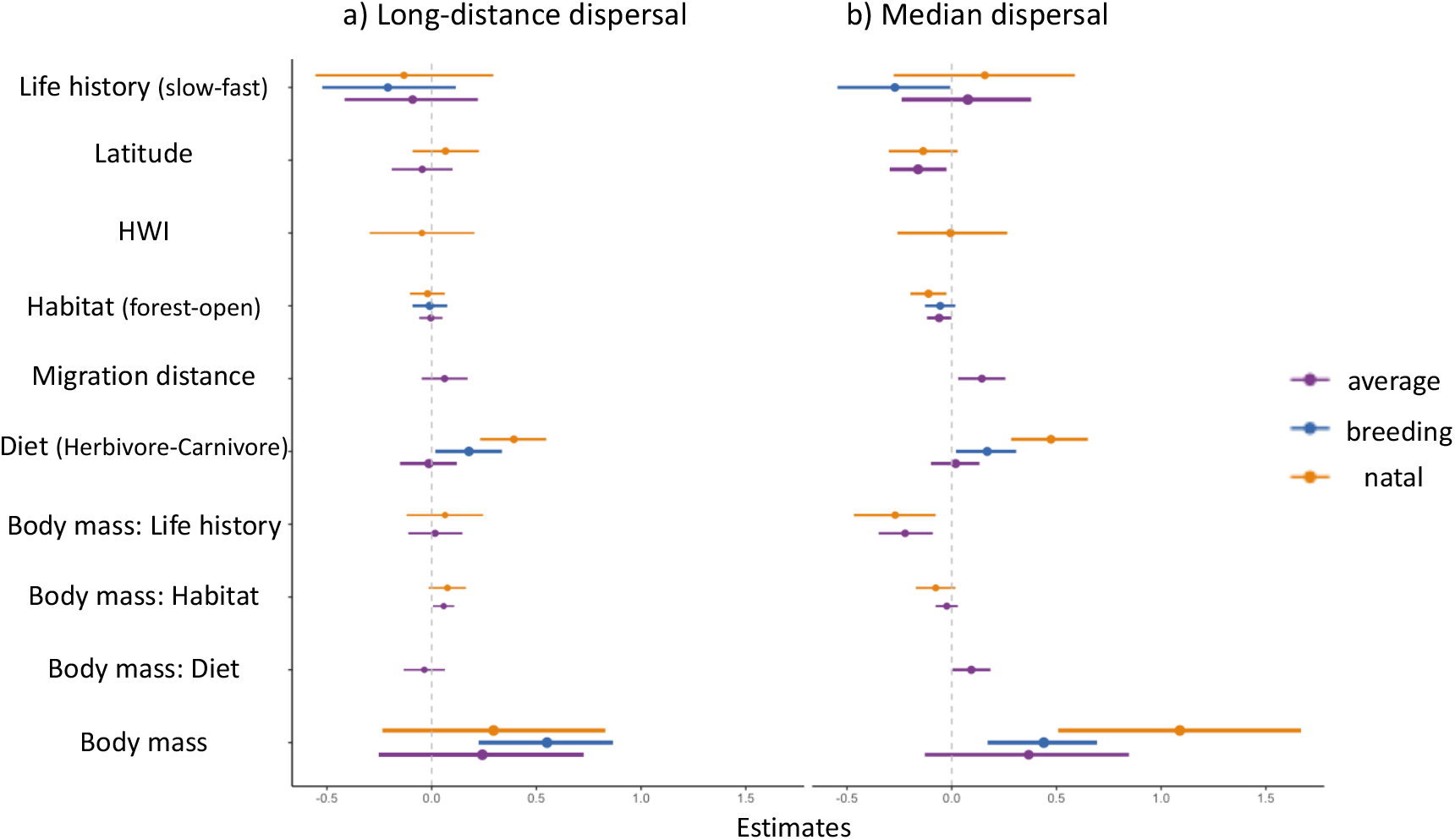
Dispersal syndromes in European birds. Standardized coefficients and 95% confidence intervals of predictors of average, breeding and natal (a) long-distance, (b) median dispersal distances among European birds based on multiple generalized linear mixed models accounting for phylogenetic relatedness. Variable importance based on mean log-predictive density (elpd) is indicated by the size of the dots. Dispersal estimates stem from the Weibull distribution(8).

**Figure 2.**
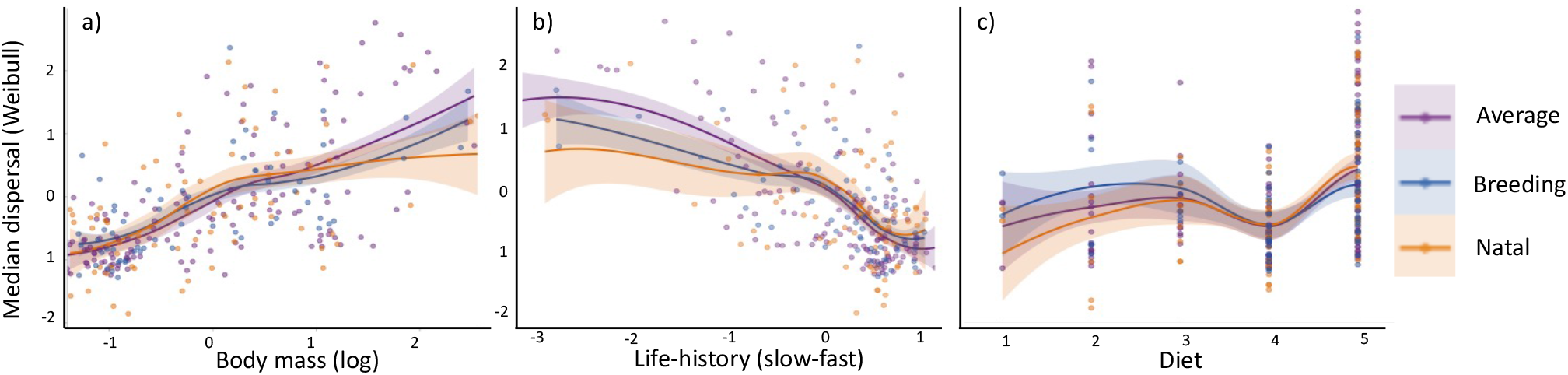
Effect of morphological, ecological, and life history traits on dispersal. Panels show how dispersal in European birds varies with (a) body mass, (b) life history traits, and (c) feeding guild. The points represent the species-specific predictions from the multi-trait model (including interactions), lines show a loess regression through the fitted point cloud and shading shows 95% confidence intervals. Dispersal estimates stem from the Weibull distribution ^8^.

### Predicting dispersal within and across orders

We assessed the predictive accuracy of single-trait and multi-trait models to estimate dispersal for missing species within and between orders. For the within-order cross-validation, we used a five-fold design refitting the trait models to 80% of the species and cross-predicting to the hold-out 20% of the species. When considering average dispersal, single-trait models, including body mass (R^2^ = 0.454) and life history (R^2^ = 0.407), respectively, achieved the highest predictive power for the median dispersal. In comparison, the multi-trait model (i.e., the multi-trait dispersal syndrome) only achieved predictive power of R^2^ = 0.263 for the median dispersal (Fig. 3; Table S2). Thus, single-trait models achieved higher within-order predictive performance than the multi-trait model and the models only calibrated with phylogeny or randomly (Fig. S4). In accordance with these results, body mass (R^2^ = 0.546) and life history (R^2^ = 0.516) also achieved the highest predictive power for natal dispersal, while body mass (R^2^ = 0.638) and habitat used (R^2^ = 0.629) best predicted median breeding dispersal.

**Figure 3.**
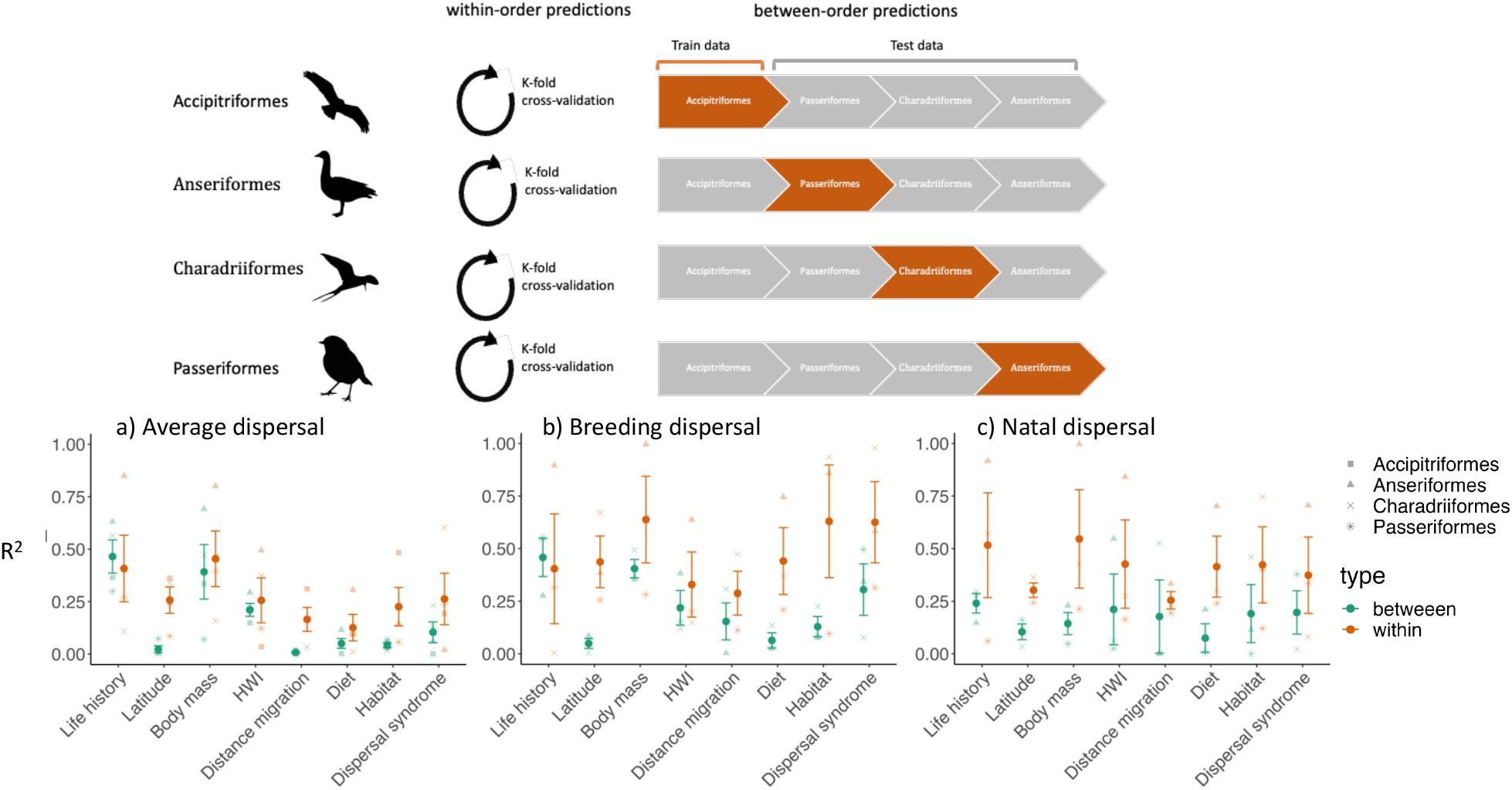
Predictive accuracy of single-trait and multi-trait dispersal models. Top: Conceptual overview of within-order and between-order cross-validation approach. We only included orders with min. 5 species for average dispersal (Accipitriformes 11 sp; Anseriformes 12 sp; Charadriiformes 11 sp; Passeriformes 68 sp), and for natal and breeding dispersal (Anseriformes 6 sp; Charadriiformes 8 sp; Passeriformes 36 sp). Bottom: Predictive performance in terms of R2 obtained from the within and between-order cross-validation for single-trait and multi-trait models (dispersal syndrome). a) average dispersal, b) natal dispersal, and c) breeding dispersal. Point shapes show the four different orders used to calibrate the models, and the colors indicate predictions within and between orders.

For the between-order cross-validation, we used a four-fold design and selected four orders with a reasonable number of species (11-68 species richness within order), refitted the trait models with one order and cross-predicted to the other three orders to assess prediction accuracy (Fig. 3; Table S2). This is a very conservative scenario reflecting that typically we have dispersal estimates for fewer species than we predict to ^10^. Again, for average dispersal, the highest between-order predictive performances were achieved in single-trait models of body mass (R2= 0.392) and of life history (R2= 0.465), and these models clearly outperformed predictions based only on phylogeny or calibrated randomly (Fig. S3). Similarly, for breeding dispersal, between-order predictions were best for the single-trait models of body mass (R2= 0.404) and life history (R2= 0.458). For natal dispersal, life history (R2= 0.240) and HWI (R2= 0.211) achieved the highest predictive power. As could be expected, within-order predictions had better predictive power than between-order predictions, especially for breeding and natal predictions (Fig. 3). Also, within-order predictions and between-order predictive performance were lower for average dispersal than for natal and breeding dispersal (Fig. 3). Long-distance dispersal presented similar patterns, but overall lower predictive accuracy in all types of dispersal, especially for between-order predictions (Fig. S2; Table S3). These results show that dispersal syndromes are valuable for understanding variation in dispersal traits of closely related species, but dispersal ability across orders can be better predicted from single traits (mainly life-history and body mass) derived from widely accessible databases. The complex underlying mechanisms and multidimensionality of dispersal, especially for long-distance dispersal, may preclude predictions using a multi-trait approach.

## Discussion

Our comprehensive analysis of dispersal syndromes in European birds revealed consistent covariations between dispersal distances and a suite of morphological, life-history and ecological traits. Importantly, although multi-trait dispersal syndromes help understand within-order variation in dispersal, between-order variation in dispersal is better predicted by single-trait models related to body mass or life history that both have clear mechanistic meaning. Thereby, median- and long-distance dispersal as well as natal vs. breeding dispersal, seem to rely on largely similar mechanisms. Our work provides unique insights into the determinants of broad-scale inter-specific variation in dispersal ability and important implications for predicting dispersal for species missing empirical estimates ^22,35,36^.

Dispersal syndromes serve as valuable tools in studying and comprehending the diverse patterns of dispersal observed among different species. Our interspecific comparisons show that multiple selection pressures are likely to play a role in driving species dispersal ability. The results support predictions from theoretical models and empirical work that have suggested that species with larger body sizes and high fecundity display higher dispersal ability ^15,22,26,36–38^. This can be attributed to factors such as the size-dependent nature of competitive ability, energy availability, and energetic requirements and how dispersal reduces competition and avoids inbreeding ^38,39^. Diet is another factor partially associated with dispersal. We found that carnivorous birds exhibit higher dispersal tendencies than other trophic groups. This is likely due to their need to cover larger areas in search of suitable prey and to optimize foraging success, resulting in increased dispersal distances ^20,38,39^. We found interactions between several traits and body mass determined bird dispersal, suggesting that competing mechanisms are linked to bird size. Surprisingly, other traits analyzed (e.g. migratory distance, habitat or latitude) were only weakly associated with dispersal distances ^15,38,40^. Although previous comparative studies show that HWI and wing morphology are strong predictors of dispersal ^16,17,41,42^, we found minimal or negligible influence of HWI on natal, breeding and average dispersal distances. This discrepancy may arise from the inherent challenges of disentangling migration and movement capacity from dispersal behavior ^43–45^. Further research is needed to elucidate the mechanisms underlying dispersal decisions and the potential interplay between movement capacity, migration, and dispersal behavior in avian populations.

Measuring bird dispersal usually requires extensive mark-recapture studies or direct tracking of individuals over multiple years, which is always costly and may prove impossible ^9^. We provide evidence that the trait-based approach can be useful for inferring median and long-distance dispersal distance, but our results also shed light on the challenges and limitations associated with predicting dispersal in species without dispersal information ^6,25^. Considering phylogenetic relatedness is certainly important when predicting ecological traits and behaviors ^46^. The shared evolutionary history within taxonomic orders facilitates understanding trait covariation and improves predictions within these groups. However, predicting dispersal across taxonomically distant orders with a multi-trait model is challenging due to the wide trait variation, introducing complexity and uncertainty with very limited predictive power at low sample sizes ^47^. Predictions from single traits such as body size and life history that have clear mechanistic relationships with dispersal ^15,25,38^ seem much more robust when predicting across orders (while controlling for phylogenetic relatedness). In contrast, the multi-trait approach, which encompasses traits such as habitat, migration distance and diet, may have limited predictive power due to their higher trait variability across the phylogenetic tree^48,49^. This emphasizes the complexity of dispersal and the different selective pressures and phylogenetic pathways among different clades ^14^. Regardless of the underlying mechanisms, the lower between-order predictive performance observed for natal dispersal and for long-distance dispersal suggests that the lack of prior knowledge regarding resource distribution and mating opportunities in unfamiliar territories makes these dispersal movements more context-dependent and influenced by environmental conditions ^39,50^.

Although representing the, to date, most comprehensive study on dispersal across the tree of life, several outstanding questions remain. First, our analyses were limited to European birds, which may restrict the generalizability of our results to other biomes and taxa with different ecological and evolutionary dynamics. However, extending the scope of investigations to other geographic regions and taxa will only be possible if more comprehensive dispersal databases become available ^8^. Second, dispersal kernels are phenomenological and context-dependent emerging from different underlying mechanisms and drivers ^7,51^. Dispersal is a complex process involving distinct phases, including departure, transfer, and settlement, each influenced by many factors operating across various spatial and temporal scales that may impact dispersal syndromes or our dispersal ability predictions ^9,52^. We conducted the analysis at the species level and hence did not consider intra-specific variation in the dispersal syndrome and context dependence ^50^. Future research should assess the consistency of dispersal syndromes at the intra-specific level ^14,50,53,54^ and should complement phenomenological dispersal kernels with mechanistic models incorporating individual behavior, environmental factors, and evolutionary processes ^55^.

Dispersal distance is an essential input for process-based population models ^11^, that offer more robust predictions of movements and interactions between species, species ability to adapt to changing environments, and ultimately develop effective strategies for conservation and management in the face of ongoing environmental challenges ^11,56,57^. Our research underlines the existence and multi-faceted nature of dispersal syndromes, emphasizing the need for a more comprehensive framework to predict dispersal and its implications in changing environments across diverse taxa and ecosystems.

## Materials and Methods

### Dispersal data

We gathered standardized estimates of median and long dispersal from ^8^.These are based on mark and recapture data from the EURING database ^58^ and provide average dispersal kernels (pooling all age classes) for 234 species, breeding dispersal kernel (between subsequent breeding attempts) for 113 species, and natal dispersal kernels (from natal site to first breeding site) for 122 species of European birds. We used the dispersal estimates from the Weibull and Half-Cauchy distributions because these clearly outperformed others for capturing rare long-distance dispersal events that contribute significantly to population dynamics ^8^. In subsequent analyses, we used both the empirical median dispersal distance from the dispersal kernels and the long-distance dispersal measures, which were defined as the 95% percentile of the dispersal kernel.

### Traits

Species’ traits were represented by 7 variables related to morphology, life history, diet, behavior, habitat and range geography (Table S1): (a) body mass, (b) Hand Wing Index, (c) diet niche, (d) life history strategy, (e) habitat preference in terms of landscape openness, (f) mean latitude of breeding range, and (g) migration distance between summer breeding and winter range. Body mass and Hand Wing Index (HWI), measured as Kipp’s distance corrected for wing size, were extracted from ^17^. Diet niche position was defined by an ordinal measure ranging from species feeding obligatory on plants (1) to species feeding obligatory on animals (5) ^31^. Life history strategy was defined as the position of species along the slow-fast life history axis obtained by a principal component analysis of five life history traits extracted from ^32^, namely egg mass, clutch size, age of first breeding, number of broods per season and life span. The first axis of the principal component analysis represents a gradient between so-called “K-selected” species with slow strategies and “r-selected” species with fast strategies. Habitat openness preference was defined as position of species’ habitat niche along the gradient from forest interior (value = 1) to open treeless landscape (7) ^59^. Finally, the mean latitude of breeding range was extracted from ^59^ and the migration distance was measured as a great circle distance between centroids of species’ breeding and non-breeding ranges ^60^. Before subsequent analyses, all traits were scaled and standardized by subtracting the mean and dividing by the standard deviation ^61^. Variance inflation factors (VIF) between all traits used in the analysis were always below 5. Because trait data were unavailable for all species in our data set, the number of species with the complete dataset was 138 species for average, 63 for breeding and 72 species for natal dispersal. We tried to develop a dataset with phylogenetic trait imputation to fill gaps in our data set. However, error estimations had high variability, and results were sensitive to the trait imputation (Supplemental material), making it difficult to disentangle if results differed because of the error estimation from trait imputation or because new species were included. Thus, we decided for the subsequent analysis to continue with only complete datasets (see Supplementary material for trait imputation results).

### Statistical analyses

#### Quantifying dispersal syndromes

We used Bayesian phylogenetic mixed models implemented in the package “brms” ^62^ to analyze the relationships between species traits and dispersal. We accounted for phylogenetic relationships among species by including a covariance matrix containing phylogenetic distances among species. We obtained a species-level phylogeny on a sample of 500 trees obtained from the Hackett backbone of the global bird phylogeny (www.birdtree.org) ^30^, and the phylogenetic tree was transformed into a variance–covariance matrix ^47^ using the vcv.phylo() function in the package “PHYTOOLS” ^63^. We used a variance inflation factor (VIF) analysis to account for potential multicollinearity, and all traits were retained in the model if VIF <5. For each model, we obtained posterior distributions of all parameters by running 4 chains in parallel for 1,000 iterations discarding the first 500 as burn-in. We used the function get_prior() in “bmrs” package to set uninformative, flat priors for the fixed effects ^62^. Convergence was assessed by visually inspecting trace plots and ensuring that the R-hat parameter was 1 or close to 1 (≤1.02 in all cases). We report posterior means and their 95% credible intervals (CI) for all effects. Model r-squared values were computed using function r2_bayes() from the package “performance” ^64^.

We ran models for each dispersal descriptor (median and long-distance dispersal), for each dispersal type (average, breeding and natal) and for the two main dispersal kernel distributions (Half-Cauchy and Weibull; ^8^. In each case, we fitted three models, one null model with no covariates (only phylogeny), one full model with all traits, and one variable-selected multi-trait model based on forward stepwise variable selection. We used the projection predictive inference for the selection of relevant predictor traits because it provides an excellent trade-off between model complexity and accuracy ^65^, especially when the goal is to identify a minimal subset of traits that yield a good predictive model.

The projection predictive inference approach can be viewed as a two-step process. First, we fitted the full model with all traits. Note that this full model only included main effects and the interactions between body size and life history, diet, and habitat openness. In the second step, the goal is to replace the posterior distribution of the reference model with a simpler distribution. This is achieved via a forward step-wise addition of traits that decrease the Kullback–Leibler divergence of the model compared to the reference (full) model. By inspecting the mean log-predictive density (elpd) and root-mean-squared error (r.m.s.e.) for each number of traits, we chose the model with the smallest number of traits yet similar predictive performance as the full model (in terms of elpd). The stepwise variable selection was implemented using the R package “projpred” ^66^. To get a useful predictor ranking, we manually delayed the phylogenetic covariance matrix term to the last position in the predictor selection process. Without this, it would take in almost all the variance of the dependent variable. In all cases, we compared the reduced model with the selected variables, the full and null model based on elpd using the “loo” package ^67^.

#### Evaluating and cross-predicting dispersal estimates

We tested the predictive performance of dispersal syndromes within and between bird orders, comparing our variable-selected multi-trait models and single-trait models. This way, we were able to ascertain how generalizable predictions of dispersal distances are across the bird phylogenetic tree when based on dispersal syndromes, including multiple traits or single traits. We selected four bird orders with a reasonable number of species to test within-order and between-order predictive performance (*Accipitriformes* 11 species, *Anseriformes* 12 species, *Charadriiformes* 11 species and *Passeriformes* 68 species) and used dispersal distances from the Weibull distribution ^8^.

Within-order predictive performance was assessed using five-fold cross-validation where species were partitioned into five folds, the multi-trait or single-trait models retrained on four folds and predicted to the hold-out fold of species ^68^. Between-order predictive performance was assessed by training the multi-trait and single-trait models on each of the four bird orders and then predicting dispersal distances to the remaining orders. In all calibrated models, we included the covariance matrix containing phylogenetic distances among species. In the test, we allowed the predictions the possibility of including new levels on this covariance matrix, meaning that the prediction will use the unconditional values for data with previously unobserved levels. To examine the predictive power of the single-trait and multi-trait models, and the ability of models to predict dispersal distances between and within orders correctly, we used the function model_perfomance() from “performance” R package ^64^. We used the r-squared value to evaluate the predictive performance ^69^.

## Author Contributions

GF. and DZ. designed research; GF., performed research; GF. analyzed data; GF., and DZ. conception and management of the project; GF wrote the first draft; and GF., RR., and DZ. write the paper.

## Code availability

The code used to analyse the data and create the figures in this paper are available in Zeonodo, with the identifier: Guillermo Fandos, Rob, R., & Damaris, Z. (2024). Dispersal syndromes on European Birds (submitting). Zenodo. https://doi.org/10.5281/zenodo.10713958.

## Acknowledgments

DZ and GF received funding from the German Science Foundation DFG (grant no. ZU 361/1-1)

