## Supplemental Material for "Dispersal syndromes allow understanding but not predicting dispersal ability across the tree of life"

**Supplementary Information:**

### Trait data imputation

Because trait data (Table S1) were not available for all species in our data set, we used phylogenetic trait imputation to fill gaps in our data set. Trait imputation can provide reliable information for up to 60% of missing data, and adding phylogenetic information to trait imputation has been shown to strongly reduce estimation error [1]. However, estimation error may be higher when closely related species have missing data. We used a random forest imputation algorithm in combination with phylogenetic information with the missForest function in the R package missForest [2]. For each trait, we assessed imputation error for a range of phylogenetic eigenvectors (1–30) and used the imputed trait values for the number of phylogenetic eigenvectors that minimized imputation error. Our gap-filled trait data had similar distributions and median values as the observed trait data. However, results were sensitive to the imputed trait data. Thus, we decided to remove species with missing values from the analysis, yet also present results excluding species with imputed trait data. The complete datasets with all traits were 138 species for average dispersal, 63 for breeding dispersal and 72 species for natal dispersal.

### Sensitivity analysis on predictions

We tested the predictive performance of dispersal syndromes within and between bird orders, comparing our variable-selected multi-trait models and single-trait models with a model only calibrated with the phylogeny and a random null model (only intercept; Fig S3). This way, we could ascertain how robust our predictions of dispersal distances are across the bird phylogenetic tree. We selected four bird orders with a reasonable number of species to test between and within-order predictive performance (Accipitriformes 11 species, Anseriformes 12 species, Charadriiformes 11 species and Passeriformes 68 species).

Within-order predictive performance was assessed using five-fold cross-validation where species were partitioned into five folds, the multi-trait or single-trait models retrained on four folds and predicted to the hold-out fold of species (Fig S3). Between-order predictive performance was assessed by training the multi-trait and single-trait models on each of the four bird orders and then predicted dispersal distances to the remaining (Fig S3). We included the covariance matrix containing phylogenetic distances among species in all calibrated models, except for the random null model. In the test, we allowed the predictions the possibility of including new levels on this covariance matrix, meaning that the prediction will use the unconditional values for data with previously unobserved levels. To examine the predictive power of the single-trait, multi-trait models, only phylogenetic model, and random null model, and the ability of models to predict dispersal distances between and within orders correctly, we used the function 'model_perfomance' from performance R package [3]. We used the r-squared value to evaluate the predictive performance.

### Supplementary Figures and Tables

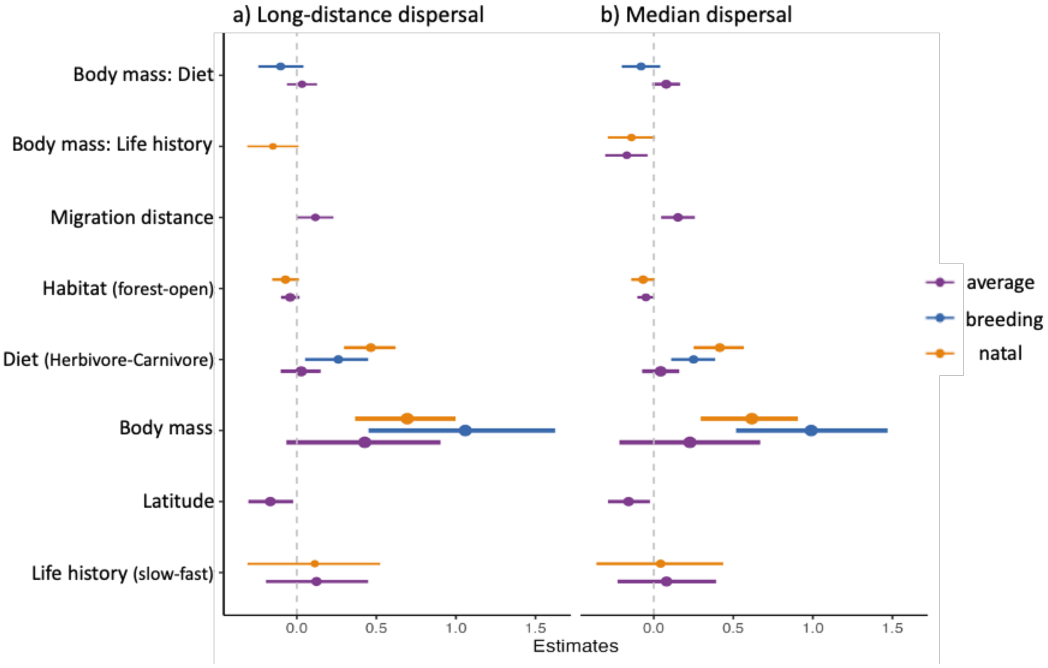

**Fig. S1. Dispersal syndromes in European birds.** Standardized coefficients and 95% confidence intervals of predictors of average, breeding and natal (a) long-distance, (b) median dispersal distances among European birds based on multiple generalized linear mixed models accounting for phylogenetic relatedness. Variable importance based on mean log-predictive density (elpd) is indicated by the size of the dots. Dispersal estimates stem from the Half-Cauchy distribution [4].

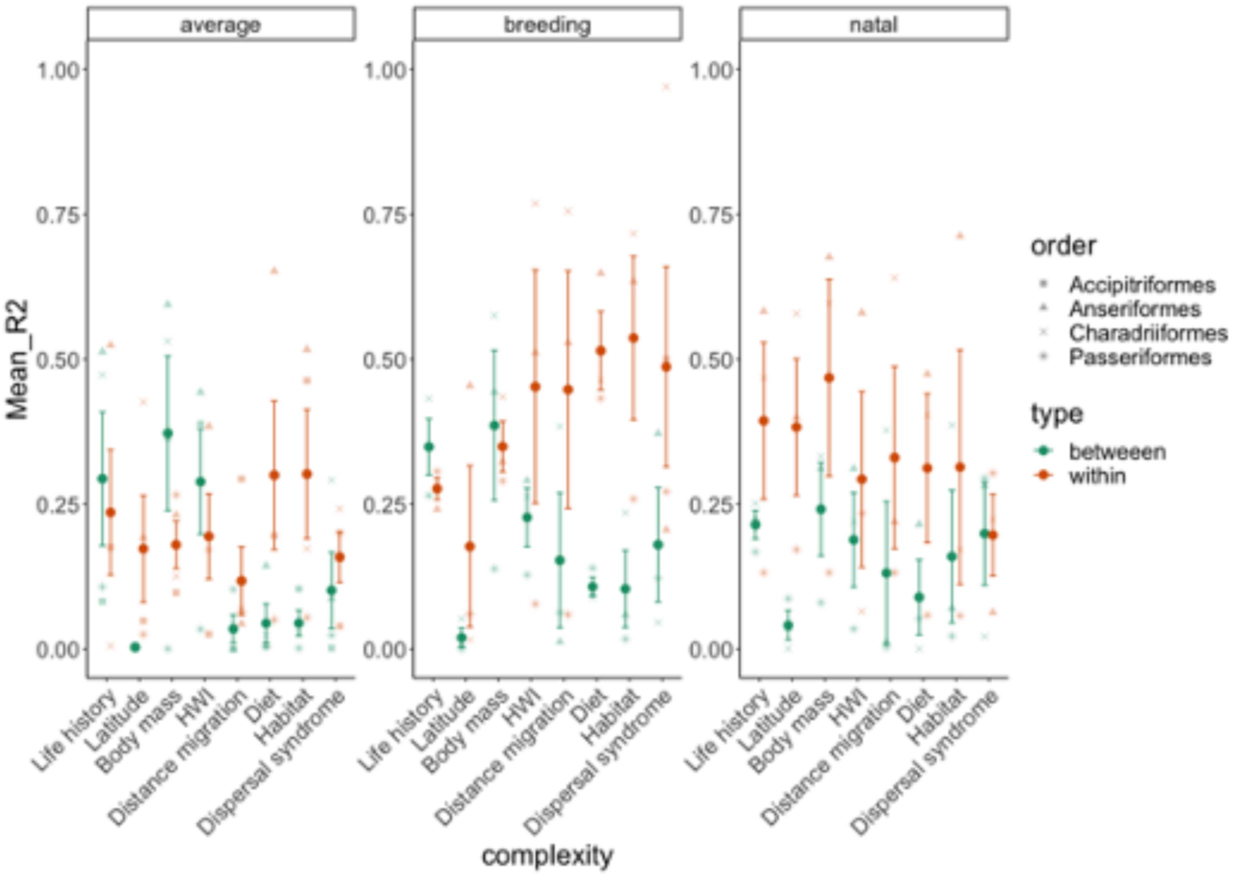

**Fig. S2.** Predictive accuracy (R2) from the within-order and between-order cross-validation for single-trait models (univariate models) and the multi-trait model (dispersal syndrome) for long-distance dispersal. a) average dispersal, b) natal dispersal, and c) breeding dispersal.

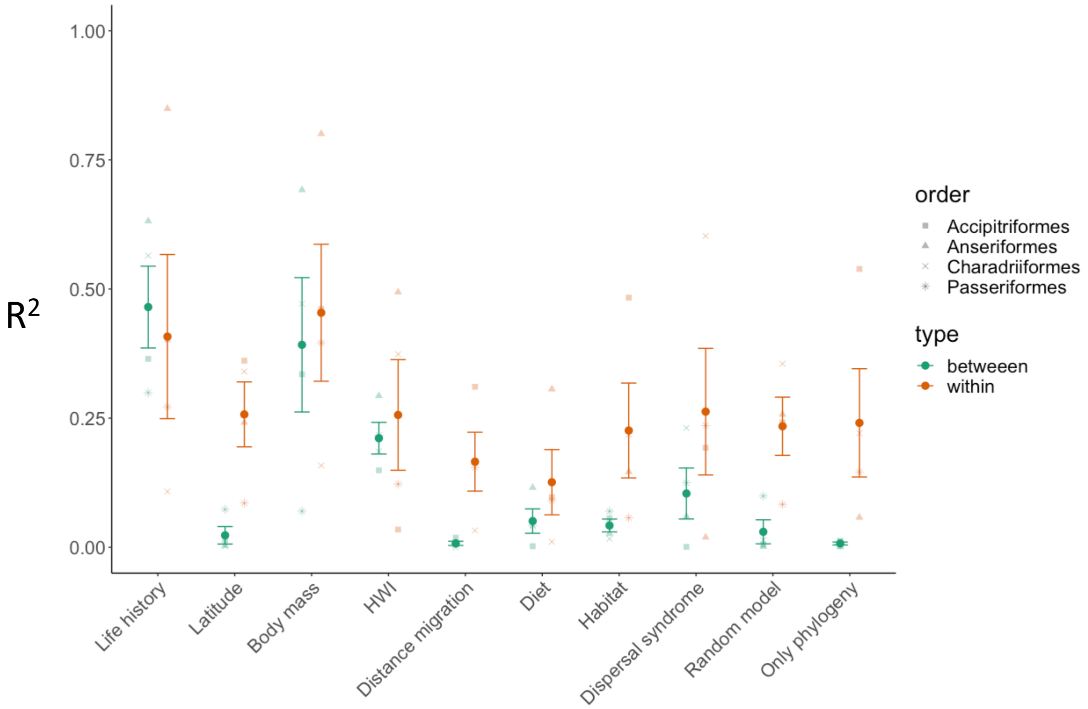

**Fig. S3**. Predictive performance (R2) from the within-order and between-order cross-validation for single-trait models (univariate models), random model (intercept only), phylogenetic model (only with a covariance matrix containing phylogenetic distances among species), and the median-distance multi-trait model (dispersal syndrome). Random model: within (R2= 0.234, sd= 0.113) and between predictions (R2= 0.030, sd= 0.046). Phylogenetic model : within (R2= 0.241, sd= 0.209) and between predictions (R2= 0.007, sd= 0.005).

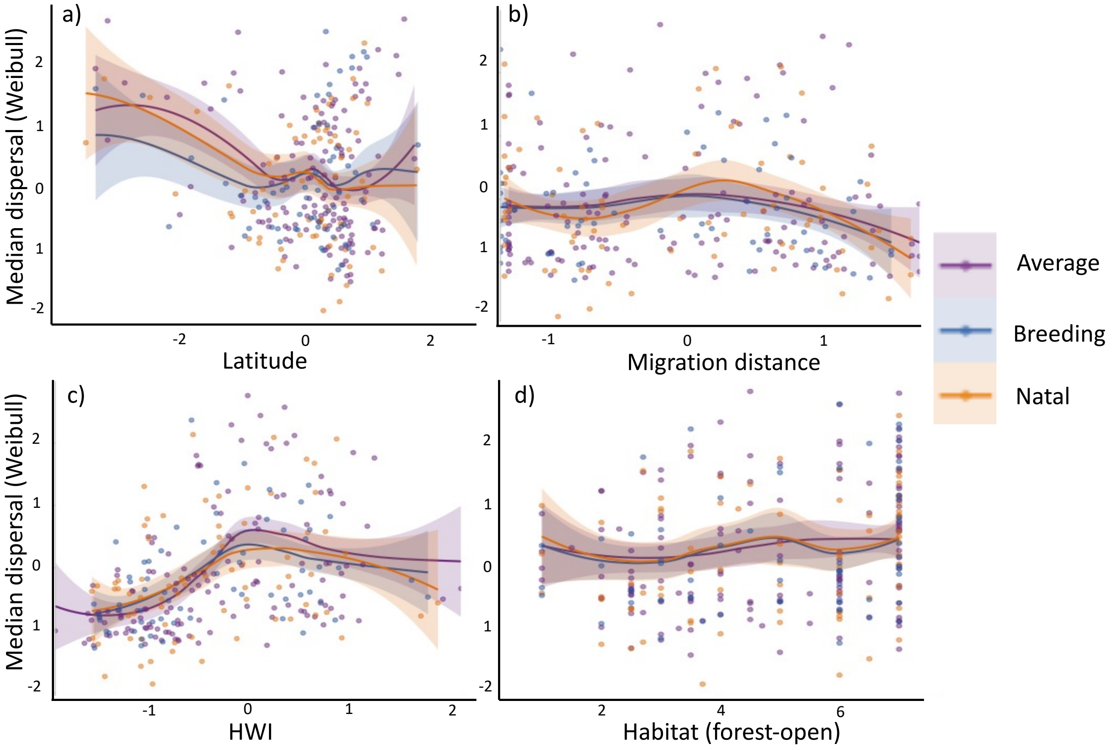

**Fig. S4.** Panels show how dispersal of European birds varies with (a) Latitude, (b) migration distance, (c) HWI, and d) habitat. The line shows a loess regression through the fitted point cloud, and shading shows 95% confidence intervals. Dispersal estimates stem from the Weibull distribution[4].

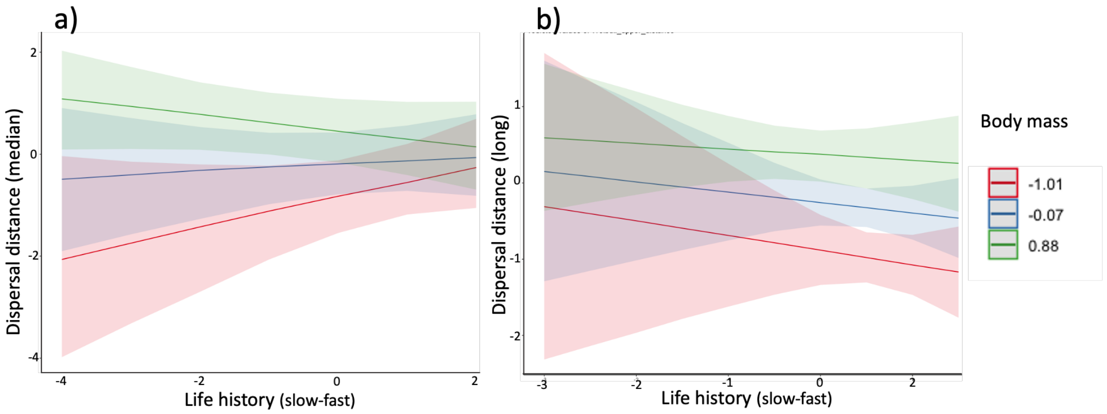

**Fig. S5**. Panels show how predicted values for a) median and b) long-distance dispersal of European birds vary within the life-history continuum depending on body mass. The line shows the marginal effects of a two-way interaction between life history and body mass and shading shows 95% confidence intervals. Dispersal estimates stem from the Weibull distribution[4]

**Table S1.** Characterization of the traits considered for analyzing European bird dispersal syndromes.

| *Trait ID* | *Trait name* | *Description* | *References* |
|---|---|---|---|
| *a* | Body mass | body weight (mean for male and female in breeding season) | Storchova and Horak (2018) |
| *b* | Hand Wing Index [HWI] | Kipp's distance corrected for wing size | Sheard et al. 2020 |
| *c* | Diet | diet niche position along the gradient from species feeding obligatory on plants (1) to species feeding obligatory on animals (5) | Reif et al. 2016 |
| *d* | Life history strategy | position along the slow-fast life history gradient revealed by a principal component analysis on six species' life-history traits | Storchova and Horak (2018); Hanzelka et al. 2019 |
| *e* | Habitat preference | mean of species' positions along the gradient from closed forest (1) to open treeless habitat (7) | Koschova et al. 2018 |
| *f* | Latitude | latitude of the center of species' breeding range in Europe (in decimal degrees) | Koschova et al. 2019 |
| *g* | Migration distance | migration distance as a Great circle distance between centroids of species' breeding and non-breeding ranges | Hanzelka et al. 2019 |

**Table S2.** Predictive accuracy of single-trait and multi-trait models to estimate median dispersal for missing species within and between orders for natal, breeding, and average dispersal. For the within-order cross-validation, we used a five-fold design refitting the trait models to 80% of the species and cross-predicting to the hold-out 20% of the species. For the between-order cross-validation, we used a four-fold design and selected four orders with a reasonable number of species (Accipitriformes 11 species, Anseriformes 12 species, Charadriiformes 11 species and Passeriformes 68 species), refitted the trait models with one order, and cross-predicted to the other three orders to assess prediction accuracy.

| Dispersal type | Prediction | Complexity | Mean $R^2$ | sd | se | ci |
|---|---|---|---|---|---|---|
| **average** | betweeen | Life history | 0,465 | 0,158 | 0,079 | 0,252 |
| average | betweeen | Latitude | 0,023 | 0,034 | 0,017 | 0,054 |
| average | betweeen | Body mass | 0,392 | 0,260 | 0,130 | 0,414 |
| average | betweeen | HWI | 0,211 | 0,061 | 0,031 | 0,098 |
| average | betweeen | Distance migration | 0,008 | 0,008 | 0,004 | 0,013 |
| average | betweeen | Diet | 0,051 | 0,047 | 0,024 | 0,075 |
| average | betweeen | Habitat | 0,042 | 0,025 | 0,012 | 0,040 |
| average | betweeen | Dispersal syndrome | 0,104 | 0,099 | 0,049 | 0,157 |
| average | within | Life history | 0,408 | 0,318 | 0,159 | 0,506 |
| average | within | Latitude | 0,257 | 0,126 | 0,063 | 0,200 |
| average | within | Body mass | 0,454 | 0,265 | 0,133 | 0,422 |
| average | within | HWI | 0,256 | 0,214 | 0,107 | 0,341 |
| average | within | Distance migration | 0,166 | 0,114 | 0,057 | 0,181 |
| average | within | Diet | 0,126 | 0,126 | 0,063 | 0,201 |
| average | within | Habitat | 0,226 | 0,184 | 0,092 | 0,292 |
| average | within | Dispersal syndrome | 0,263 | 0,245 | 0,123 | 0,390 |
| breeding | betweeen | Life history | 0,458 | 0,157 | 0,091 | 0,390 |
| breeding | betweeen | Latitude | 0,049 | 0,041 | 0,024 | 0,101 |
| breeding | betweeen | Body mass | 0,405 | 0,076 | 0,044 | 0,189 |
| breeding | betweeen | HWI | 0,219 | 0,143 | 0,082 | 0,355 |
| breeding | betweeen | Distance migration | 0,154 | 0,152 | 0,088 | 0,378 |
| breeding | betweeen | Diet | 0,064 | 0,062 | 0,036 | 0,155 |
| breeding | betweeen | Habitat | 0,129 | 0,084 | 0,048 | 0,208 |
| breeding | betweeen | Dispersal syndrome | 0,305 | 0,212 | 0,122 | 0,527 |
| breeding | within | Life history | 0,404 | 0,452 | 0,261 | 1,123 |
| breeding | within | Latitude | 0,437 | 0,212 | 0,123 | 0,527 |
| breeding | within | Body mass | 0,638 | 0,357 | 0,206 | 0,886 |
| breeding | within | HWI | 0,329 | 0,268 | 0,155 | 0,666 |
| breeding | within | Distance migration | 0,288 | 0,181 | 0,104 | 0,449 |
| breeding | within | Diet | 0,441 | 0,275 | 0,159 | 0,683 |
| breeding | within | Habitat | 0,630 | 0,464 | 0,268 | 1,153 |
| breeding | within | Dispersal syndrome | 0,625 | 0,334 | 0,193 | 0,830 |
| natal | betweeen | Life history | 0,240 | 0,081 | 0,047 | 0,201 |

| | | | | | | |
|---|---|---|---|---|---|---|
| natal | betweeen | Latitude | 0,105 | 0,064 | 0,037 | 0,160 |
| natal | betweeen | Body mass | 0,144 | 0,091 | 0,053 | 0,226 |
| natal | betweeen | HWI | 0,211 | 0,291 | 0,168 | 0,723 |
| natal | betweeen | Distance migration | 0,177 | 0,302 | 0,174 | 0,749 |
| natal | betweeen | Diet | 0,075 | 0,118 | 0,068 | 0,293 |
| natal | betweeen | Habitat | 0,191 | 0,239 | 0,138 | 0,594 |
| natal | betweeen | Dispersal syndrome | 0,197 | 0,179 | 0,103 | 0,444 |
| natal | within | Life history | 0,516 | 0,431 | 0,249 | 1,072 |
| natal | within | Latitude | 0,303 | 0,060 | 0,035 | 0,149 |
| natal | within | Body mass | 0,546 | 0,405 | 0,234 | 1,006 |
| natal | within | HWI | 0,426 | 0,363 | 0,210 | 0,903 |
| natal | within | Distance migration | 0,255 | 0,072 | 0,041 | 0,178 |
| natal | within | Diet | 0,414 | 0,251 | 0,145 | 0,623 |
| natal | within | Habitat | 0,423 | 0,313 | 0,181 | 0,778 |
| natal | within | Dispersal syndrome | 0,373 | 0,315 | 0,182 | 0,782 |

**Table S3.** Predictive accuracy of single-trait and multi-trait models to estimate long dispersal for missing species within and between orders for natal, breeding, and average dispersal. For the within-order cross-validation, we used a five-fold design refitting the trait models to 80% of the species and cross-predicting to the hold-out 20% of the species. For the between-order cross-validation, we used a four-fold design and selected four orders with a reasonable number of species (Accipitriformes 11 species, Anseriformes 12 species, Charadriiformes 11 species and Passeriformes 68 species), refitted the trait models with one order, and cross-predicted to the other three orders to assess prediction accuracy.

| Dispersal type | Prediction | Complexity | Mean $R^2$ | sd | se | ci |
|---|---|---|---|---|---|---|
| **average** | betweeen | Life history | 0,294 | 0,231 | 0,115 | 0,367 |
| average | betweeen | Latitude | 0,003 | 0,003 | 0,001 | 0,004 |
| average | betweeen | Body mass | 0,372 | 0,266 | 0,133 | 0,424 |
| average | betweeen | HWI | 0,289 | 0,181 | 0,091 | 0,288 |
| average | betweeen | Distance migration | 0,035 | 0,048 | 0,024 | 0,077 |
| average | betweeen | Diet | 0,044 | 0,067 | 0,033 | 0,106 |
| average | betweeen | Habitat | 0,045 | 0,043 | 0,022 | 0,069 |
| average | betweeen | Dispersal syndrome | 0,101 | 0,132 | 0,066 | 0,210 |
| average | within | Life history | 0,236 | 0,216 | 0,108 | 0,344 |
| average | within | Latitude | 0,173 | 0,184 | 0,092 | 0,293 |
| average | within | Body mass | 0,180 | 0,081 | 0,041 | 0,129 |
| average | within | HWI | 0,194 | 0,147 | 0,074 | 0,234 |
| average | within | Distance migration | 0,118 | 0,118 | 0,059 | 0,187 |
| average | within | Diet | 0,300 | 0,256 | 0,128 | 0,407 |
| average | within | Habitat | 0,302 | 0,223 | 0,112 | 0,355 |
| average | within | Dispersal syndrome | 0,159 | 0,087 | 0,044 | 0,139 |
| breeding | betweeen | Life history | 0,349 | 0,084 | 0,048 | 0,208 |
| breeding | betweeen | Latitude | 0,020 | 0,029 | 0,016 | 0,071 |
| breeding | betweeen | Body mass | 0,386 | 0,224 | 0,129 | 0,557 |
| breeding | betweeen | HWI | 0,227 | 0,087 | 0,050 | 0,216 |
| breeding | betweeen | Distance migration | 0,153 | 0,202 | 0,116 | 0,501 |
| breeding | betweeen | Diet | 0,108 | 0,028 | 0,016 | 0,070 |
| breeding | betweeen | Habitat | 0,104 | 0,116 | 0,067 | 0,287 |
| breeding | betweeen | Dispersal syndrome | 0,180 | 0,170 | 0,098 | 0,423 |
| breeding | within | Life history | 0,276 | 0,033 | 0,019 | 0,081 |
| breeding | within | Latitude | 0,177 | 0,241 | 0,139 | 0,599 |
| breeding | within | Body mass | 0,349 | 0,076 | 0,044 | 0,189 |
| breeding | within | HWI | 0,452 | 0,349 | 0,202 | 0,867 |
| breeding | within | Distance migration | 0,448 | 0,355 | 0,205 | 0,882 |
| breeding | within | Diet | 0,515 | 0,117 | 0,067 | 0,290 |
| breeding | within | Habitat | 0,537 | 0,244 | 0,141 | 0,606 |
| breeding | within | Dispersal syndrome | 0,487 | 0,346 | 0,173 | 0,550 |
| natal | betweeen | Life history | 0,214 | 0,043 | 0,025 | 0,106 |

| | | | | | | |
|---|---|---|---|---|---|---|
| natal | betweeen | Latitude | 0,041 | 0,043 | 0,025 | 0,108 |
| natal | betweeen | Body mass | 0,241 | 0,140 | 0,081 | 0,347 |
| natal | betweeen | HWI | 0,188 | 0,141 | 0,081 | 0,350 |
| natal | betweeen | Distance migration | 0,131 | 0,213 | 0,123 | 0,530 |
| natal | betweeen | Diet | 0,089 | 0,112 | 0,065 | 0,278 |
| natal | betweeen | Habitat | 0,159 | 0,198 | 0,114 | 0,492 |
| natal | betweeen | Dispersal syndrome | 0,199 | 0,154 | 0,089 | 0,381 |
| natal | within | Life history | 0,394 | 0,234 | 0,135 | 0,582 |
| natal | within | Latitude | 0,383 | 0,204 | 0,118 | 0,507 |
| natal | within | Body mass | 0,468 | 0,294 | 0,170 | 0,730 |
| natal | within | HWI | 0,293 | 0,263 | 0,152 | 0,652 |
| natal | within | Distance migration | 0,330 | 0,272 | 0,157 | 0,675 |
| natal | within | Diet | 0,312 | 0,222 | 0,128 | 0,553 |
| natal | within | Habitat | 0,314 | 0,350 | 0,202 | 0,869 |
| natal | within | Dispersal syndrome | 0,197 | 0,122 | 0,071 | 0,303 |
